# Interneurons of fan-shaped body promote arousal in *Drosophila*

**DOI:** 10.1101/2022.05.15.492037

**Authors:** Yoshiaki S. Kato, Jun Tomita, Kazuhiko Kume

**Affiliations:** Department of Neuropharmacology, Graduate School of Pharmaceutical Sciences, Nagoya City University, Nagoya 467-8603, Japan

**Author notes:** Correspondence should be addressed to, Tanabe 3-1, Mizuho, Nagoya 467-8603 Japan.

**Keywords:** sleep, arousal, circuit, dopamine, acetylcholine, *Drosophila*

## Abstract

Sleep is required to maintain physiological functions and is widely conserved across species. To understand the sleep-regulatory mechanisms, sleep-regulating genes and neuronal circuits are studied in various animal species. In the sleep-regulatory neuronal circuits in *Drosophila melanogaster*, the dorsal fan-shaped body (dFB) is a major sleep-promoting region. However, other sleep-regulating neuronal circuits were not well identified. We recently found a novel sleep-regulatory circuit consisting of arousal-promoting T1 dopamine neurons and protocerebral bridge (PB) neurons innervating the ventral part of the FB, which we named “the PB-FB pathway”. However, the post-synaptic target of the PB-FB pathway was still unknown. To identify it, we performed anterograde tracing, immunohistochemistry, and Ca^2+^ imaging analysis and found that the PB-FB pathway projects to FB interneurons, also known as pontine neurons. Besides, we found that cholinergic pontine neurons promote arousal. Moreover, we indicated that pontine neurons form an anatomical connection with sleep-promoting dFB neurons. Together, we showed that pontine neurons receive excitatory signals from the PB-FB pathway and cholinergic pontine neurons promote arousal. These results completed one of the output pathways from the PB-FB pathway.

## Introduction

Sleep is essential for many physiological functions and is conserved across mammals and invertebrates. Although sleep plays an important role in our lives, sleep control mechanisms have not been completely elucidated. To understand the mechanisms of sleep regulation, it is critical to unravel sleep-regulatory genes and neuronal circuits. *Drosophila melanogaster* has been widely used to study the mechanisms of sleep regulation^1,2^. A large number of sleep-regulatory genes have been reported in *Drosophila*^3–5^. However, the specific brain regions where those genes function to regulate sleep are not well identified. This problem makes us difficult to study molecular mechanisms of sleep regulation in detail. This problem is due to the lack of knowledge about sleep-regulatory circuits.

Regarding sleep-regulatory circuits, the central complex is of particular interest. The central complex is divided into four regions: the protocerebral bridge (PB), the fan-shaped body (FB), the ellipsoid body (EB), and the noduli (NO). In particular, the dorsal FB (dFB) neurons promote sleep^6^, the EB R5 neurons regulate sleep homeostasis^7^, and these neurons interact with each other^8^. In previous studies, we found that dopamine is a major regulator of arousal and identified a dopamine pathway from PPM3 to FB^9,10^. Recently, we found that T1 dopaminergic neurons (wakefulness-promoting), PB interneurons (sleep-promoting), and P-FN neurons (wakefulness-promoting) that project from the PB to the ventral FB and the NO form a sleep regulatory circuit, hereafter referred to as “the PB-FB pathway”^11^. However, the post-synaptic partner of the PB-FB pathway remained unclear.

In this study, we aimed to investigate the post-synaptic neurons of P-FN neurons and focused on FB interneurons, also known as pontine neurons. We found that P-FN neurons form an excitatory connection with pontine neurons. Also, we revealed that cholinergic pontine neurons promote arousal. Moreover, we discovered that pontine neurons form an anatomical connection with dFB neurons. This study provides a novel sleep-regulatory pathway that projects from the PB-FB pathway to dFB neurons.

## Materials and Methods

### Fly strains and rearing conditions

Fruit flies (*Drosophila melanogaster*) were raised at 25 °C in 50-60 % relative humidity on standard medium containing cornmeal, yeast, glucose, wheat germ, and agar, as described before^9^. They were maintained under a 12-h light: dark (LD) cycle. In this study, we used *R52B10-Gal4* (38820), *R23E10-Gal4* (49032), *R23E10-LexA* (52693), *R52B10-LexA* (52826), *UAS-mCD8::GFP* (5130), *tub-Gal80*^*ts*^ (7019), *UAS-GCaMP6s* (42746), *UAS-DenMark, syt*.*eGFP* (33064), *LexAop-P2X2* (76030), *UAS-GFP, QUAS-RFP; trans-Tango* (77124), *hDeltaC-Gal4* (75925), and *vDeltaB, C, D-Gal4* (93172) from the Bloomington *Drosophila* Stock Center, *UAS-mAChRB RNAi* (KK0107137) from the Vienna Drosophila Resource Center, and *NP2320-Gal4* (104157) from the *Drosophila* Genetics Resource Center. *UAS-dTrpA1*^12^ was a gift from Dr. Julie H. Simpson. *Cha*-*Gal80*^13^ was from Dr. Takaomi Sakai. *UAS-CD4*::*spGFP1-10, LexAop-CD4*::*spGFP11* were from Dr. Kristin Scott. *UAS-Kir2*.*1*^14^ was from Dr. Richard A. Baines. *R52B10-Gal4, R23E10-Gal4, NP2320-Gal4, UAS-Kir2*.*1*, and *UAS-dTrpA1* are backcrossed at least 5 times to the control strain (*w*^*1118*^). Male flies were used in all experiments.

### Locomotor activity and sleep analysis

The locomotor activity of individual flies was measured for 1-min intervals using the *Drosophila* activity monitoring system (TriKinetics, Waltham, MA, USA) as described previously ^9^. The flies were placed individually in glass tubes (length, 65 mm; interior diameter, 3 mm) containing 1 % agar and 5 % sucrose food at one end and were entrained for at least 3 days to LD conditions before changing to constant dark (DD) conditions. Activity data were collected continuously under LD and DD conditions. Because sleep in the daytime under LD conditions is partly regulated by light-induced suppression of locomotor activity^9^, results from DD conditions (day 2-4 of the DD) are mainly shown. Based on previous reports, sleep in *Drosophila* was defined as continuous immobile periods lasting 5 min or longer. The total activity counts and the total amount of sleep time in DD conditions were analyzed using Microsoft (Redmond, WA, USA) Excel-based software or R (R Core Team, 2020, https://www.r-project.org).

### Immunohistochemistry and Image acquisition

Whole-mount immunofluorescence staining of adult *Drosophila* brains (Figs. 1e, and 3c) was performed as previously described^15^. Other samples were imaged without staining. Adult fly brains were dissected in PBS and fixed in 4 % PFA in PBS for 20 min at room temperature. The brains were then washed three times in 0.3 % PBS-T for 20 min. After washing, the samples were blocked in 5 % normal goat serum (NGS) at 4 °C overnight. The next day, the NGS solution was replaced by primary antibody solution in 5 % NGS and incubated at 4 °C for 1 to 2 days. After washing three times, the samples were incubated in secondary antibody solution in 5 % NGS at 4 °C for 1 to 2 days. After washing three times, the brains were mounted using PermaFluor (Funakoshi). In the GFP reconstitution across synaptic partners (GRASP) experiment, monoclonal anti-GFP (G10362, ThermoFisher) at 1:100 dilution and anti-nc82 (Developmental Studies Hybridoma Bank, University of Iowa) at 1:100 were used as the primary antibodies. Alexa Fluor 488 goat anti-rabbit IgG (A11034, Invitrogen) and Alexa Fluor 568 goat anti-mouse IgG (A11004, Invitrogen) at 1:1000 were used as secondary antibodies. All brain tissues were imaged using a ZEISS LSM 800 confocal microscope (ZEISS).

**Figure 1.**
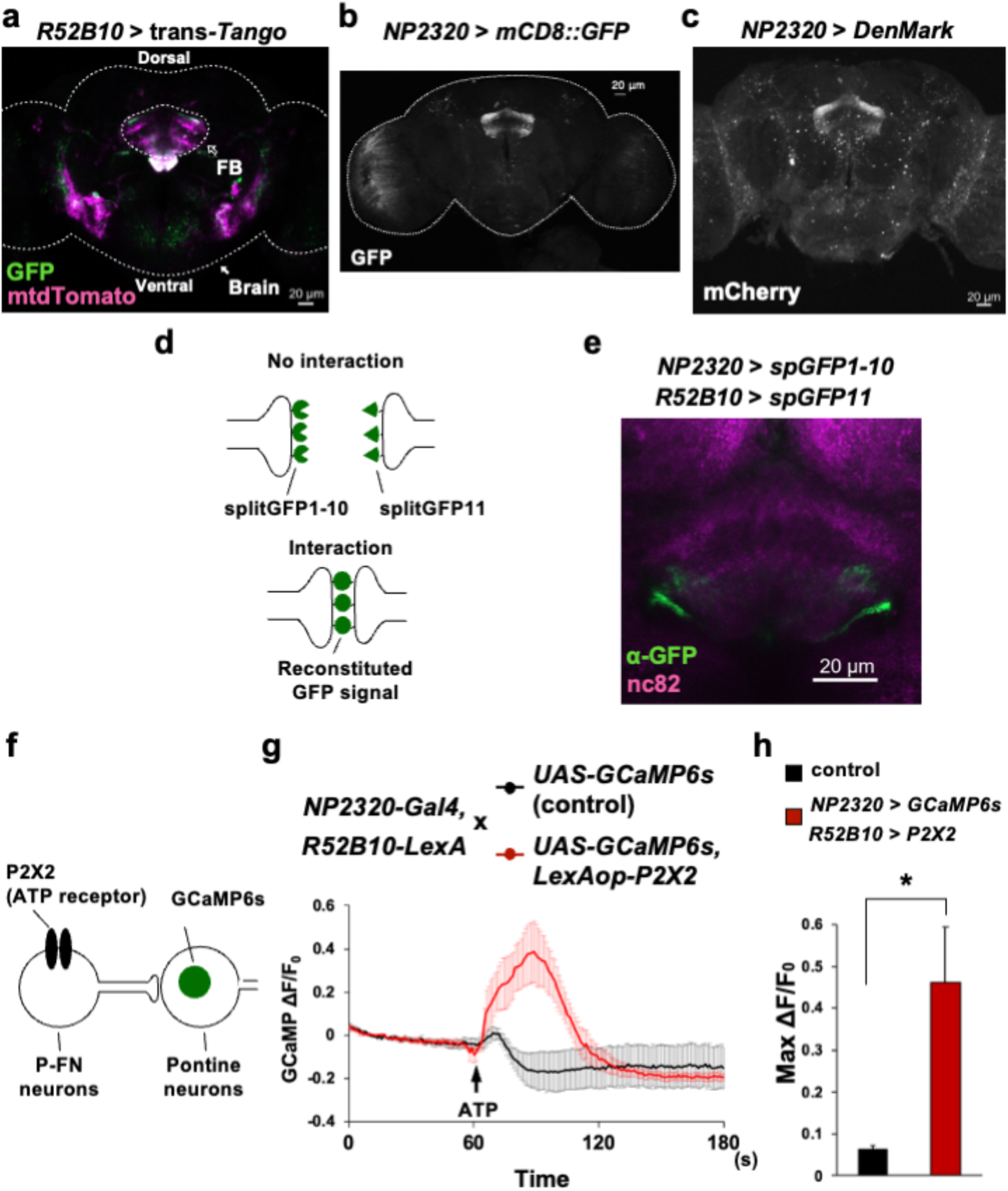
P-FN neurons and pontine neurons have an excitatory connection. **(a)** Anterograde tracing of *R52B10-Gal4*-labeled P-FN neurons using *trans*-Tango. Green shows presynaptic signals of *R52B10*-labeled P-FN neurons and magenta shows post-synaptic neuronal signals. The white arrowhead indicates the approximate outline of the brain. The black arrowhead indicates the approximate structure of the FB. The layers containing the FB are shown. **(b)** GFP signals of *NP2320-Gal4*-labeled neurons. **(c)** Expression pattern of the dendrite marker *DenMark* in pontine neurons. **(d)** Schematic representation of the GRASP method. When two neurons locate adjacent to each other, a reconstituted GFP signal is observed. **(e)** GRASP signals between *R52B10*-labeled P-FN neurons and *NP2320*-labeled pontine neurons. **(f)** Illustration of the Ca^2+^ imaging experiment. P-FN neurons that express P2X2 become activated by ATP addition and the GCaMP signals in pontine neurons are measured. **(g)** The trace of the GCaMP signals change. After adding ATP, the GCaMP signals significantly increased in the experimental group (red line, n = 5), but only slightly increased in the control group (gray line, n = 5). The y-axis indicates the change of the GCaMP signal. **(h)** Quantification of Max ΔF/F_0_. Data are presented as mean + SEM. Two-sided Welch’s *t-*test was used. * P < 0.05

### Ca^2+^ imaging

Male flies were dissected in calcium-free adult hemolymph-like saline consisting of 108 mM NaCl, 5 mM KCl, 8.2 mM MgCl_2_, 4 mM NaHCO_3_, 1 mM NaH_2_PO_4_, 5 mM trehalose, 10 mM sucrose, and 5 mM HEPES (pH 7.5). The isolated brains were placed at the bottom of a well of an 8-well plate (ibidi, Germany) beneath the adult hemolymph-like saline. All imaging was performed using a ZEISS LSM 800. To reduce the effect of the z-plane drift, the pinhole was adjusted to 105 μm. All images were taken using a 10x objective lens. Time-series images were collected for 180 s at 1 Hz. After taking baseline images for 60 s, 25 mM ATP was applied by bath application using a pipette. A region of interest (ROI) was determined based on the GCaMP baseline signal on the FB neuropile and drawn around the target structure using Fiji software (https://fiji.sc). The fluorescence signal in each ROI was analyzed using Fiji software. The transition in fluorescence was calculated following this formula: ΔF = Ft-F0/F0 (Ft: fluorescence at time point n; F0: fluorescence at time 0).

### Experimental design and statistical analysis

Data were analyzed as described in each figure Legend using Microsoft Excel and R. The number of flies used in the experiments is also described in Figure Legends.

## Results

### P-FN neurons form excitatory synaptic connections with pontine neurons

We first conducted anterograde tracing using the *trans*-Tango system to identify the post-synaptic neurons of the *R52B10-Gal4*-labeled P-FN neurons^16^. By using this system, *R52B10-*labeled P-FN neurons express GFP and their post-synaptic neurons express mtdTomato. As a result, post-synaptic signals shown in magenta were detected in the dorsal and ventral parts of the FB (Fig. 1a). Based on the morphological similarities, we assumed that one of the candidates of the post-synaptic neurons of P-FN neurons is FB interneurons, also known as pontine neurons^17,18^. Among the Gal4 drivers, *NP2320-Gal4* is a driver that is reported in the previous papers to show clear labeling in pontine neurons^19–22^. Figure 1b shows the morphological patterns of pontine neurons labeled with *NP2320-Gal4* (Fig. 1b). To examine whether pontine neurons receive signals from P-FN neurons, we labeled the dendrites of pontine neurons using the dendrite marker *DenMark*, which is mCherry-tagged hybrid protein of mammalian ICAM5/Telencephalin^23^. We found that *DenMark* was expressed in both the dFB and the vFB (Fig. 1c). This result suggests that pontine neurons have dendrites in the vFB. We then used the GRASP technique^24,25^ to confirm the connections between P-FN neurons and pontine neurons. In this technique, two different cell populations express individual split GFP components (GFP1-10 and GFP11), which reconstitute into a functional GFP molecule if these cells have close interactions (Fig. 1d). As a result, we found reconstituted GFP signals in the vFB (Fig. 1e). This result suggests that P-FN neurons and pontine neurons form synaptic connections in the vFB. To further confirm this result, we conducted ex vivo Ca^2+^ imaging. We expressed the ATP-gated cation channel *P2X2* in P-FN neurons and the Ca^2+^ indicator *GCaMP6s* in pontine neurons^26,27^ (Fig. 1f). By adding ATP to the isolated fly brain in the chamber, P-FN neurons are activated by P2X2. Then, the change in the GCaMP signals can be found if P-FN neurons and pontine neurons have functional connections. As a result, we found a substantial increase in the GCaMP signals when ATP was added to the isolated fly brain (Fig. 1g, h, and Movie S1; P = 0.038; two-sided Welch’s *t-test*). Taken together, these results suggest that P-FN neurons and pontine neurons form excitatory connections.

### Activation of pontine neurons affects sleep amounts

To investigate the role of pontine neurons in sleep regulation, we performed a transient thermo-genetic activation of pontine neurons using the thermo-activatable cation channel *dTrpA1*, which is more active at 29 °C and less active at 22 °C^12^. Flies were transferred from 22 °C to 29 °C to activate the pontine neurons and then returned to 22 °C. As a result, a significant decrease in sleep time was found on Day2 at 29 °C (Fig. 2a and S1). Moreover, *Cha-Gal80*, which inhibits Gal4 activity in the cholinergic neurons, suppressed this phenotype (Fig. 2a and S1). These results indicate that cholinergic pontine neurons promote arousal. To confirm that arousal-promoting pontine neurons are cholinergic, we investigated whether *Cha-Gal80* suppressed GFP expression in pontine neurons. From the result in Fig 2b, we found that GFP signals in pontine neurons almost disappeared compared to Fig 1b (Fig 2b). Altogether, these results indicate that cholinergic pontine neurons promote arousal.

**Figure 2.**
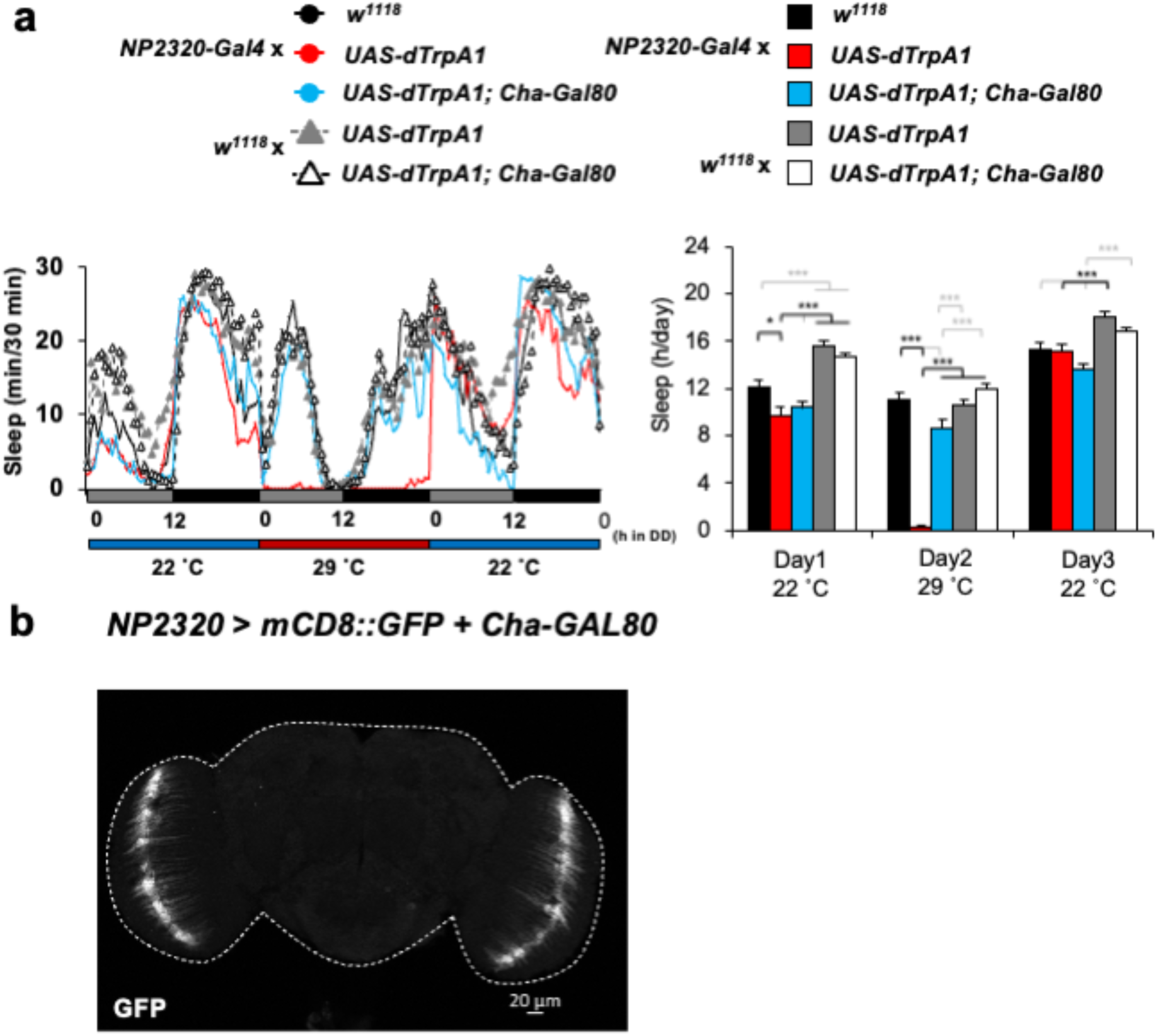
Cholinergic pontine neurons promote arousal. (a) right: Sleep profile of each genotype (n = 20, 32, 21, 32, 32 respectively). Thermo-genetic activation by *dTrpA1* occurs at 29 °C but not at 22 °C. left: Quantification of the sleep time. Data are presented as mean + SEM. one-way ANOVA with a Tukey-Kramer HSD test was used. * P < 0.05, ** P < 0.01, *** P < 0.001 (b) GFP expression pattern of *NP2320-Gal4* with *Cha-Gal80*.

**Figure 3.**
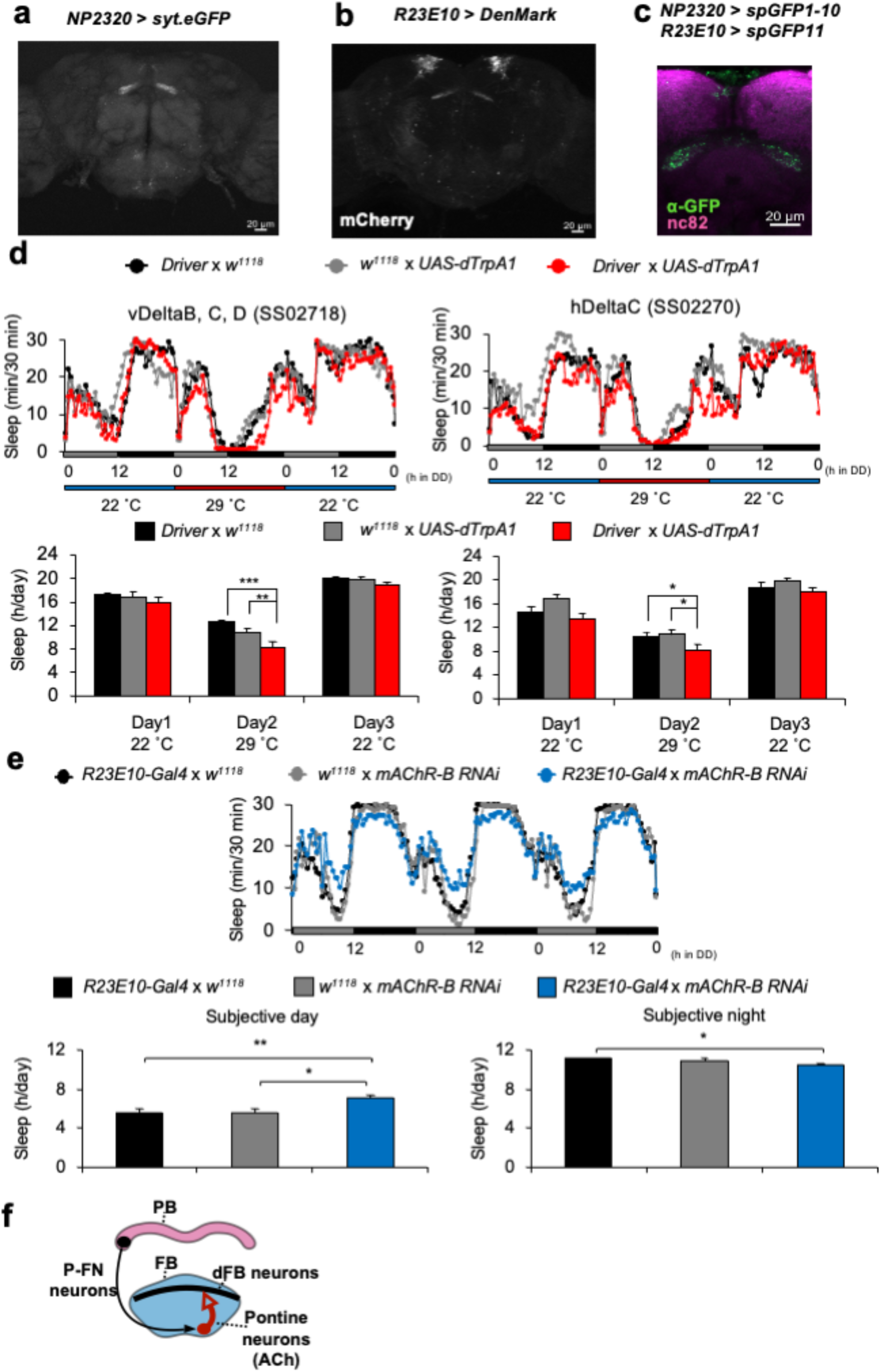
Pontine neurons send arousal signals to the dFB neurons. (a) Expression pattern of *syt*.*eGFP* in *NP2320*-labeled neurons. (b) Expression pattern of *DenMark* in *R23E10*-labeled neurons. (c) GRASP signals between *R23E10*-labeled dFB neurons and *NP2320*-labeled pontine neurons. (d) top: Sleep profile of each genotype (n = 16, 16, 16, 14, 16, 16 respectively). Thermo-genetic activation by *dTrpA1* occurs at 29 °C but not at 22 °C. bottom: Quantification of the sleep time. Data are presented as mean + SEM. one-way ANOVA with a Tukey-Kramer HSD test was used. * P < 0.05, ** P < 0.01, *** P < 0.001 (e) top: Sleep profile of each genotype (n = 32, 16, 32 respectively). bottom: Quantification of the sleep time. Data are presented as mean + SEM. one-way ANOVA with a Tukey-Kramer HSD test was used. * P < 0.05, ** P < 0.01 (f) Schematic summary of the findings. P-FN neurons activate cholinergic pontine neurons and promote arousal.

### Pontine neurons send arousal signals to dFB neurons

To investigate the post-synaptic partners of pontine neurons, we expressed *syt*.*eGFP*, an axon terminal marker, in pontine neurons. We found that the axon terminals of pontine neurons were arborized in the dorsal part of the FB (Fig. 3a). From this result, we hypothesized that one of the post-synaptic partners of pontine neurons is the sleep-promoting dFB neurons. Then we expressed *DenMark* in dFB neurons with *R23E10-Gal4*. We found the dendrites of dFB neurons also arborized in the dorsal part of the FB (Fig. 3b). We next conducted GRASP experiments to investigate the connections between pontine neurons and dFB neurons. As a result, we found GRASP positive signals in the dorsal part of the FB (Fig. 3c). These results suggest that pontine neurons and dFB neurons form anatomical connections in the dFB. Then we wondered what types of FB interneurons convey the arousal information from P-FN neurons to dFB neurons. By using the connectome database and the knowledge from the connectome paper^28^, we chose types of neurons as candidates. These are named vDeltaB, C, D, and hDeltaC neurons, which convey information from P-FN neurons to the FB layers 6 and 7. We picked up two Gal4 driver lines which show restricted expression of Gal4 in vDeltaB, C, D, and hDeltaC neurons (SS02718 and SS02270 respectively) by using the connectome database and conducted the same experiment as figure 2a. We found that the amount of sleep was decreased in both two cases (Fig. 3d). Together, we concluded that pontine neurons send arousal signals to dFB neurons. Next, we investigated the role of acetylcholine in dFB neurons regulation. Because the activation of pontine neurons decreased sleep and dFB neurons promote sleep, we focused on the inhibitory cholinergic signals. In *Drosophila melanogaster*, one inhibitory acetylcholine receptor, which is Gi-coupled muscarinic acetylcholine receptor *mAChR-B*, is reported^29,30^. We knockdown mAChR-B in the dFB neurons using R23E10-Gal4 and found that the amount of sleep was increased especially on subjective days (Fig. 3e). These results suggested the possibility that acetylcholine from pontine neurons inhibits dFB neurons via mAChR-B and promotes arousal. Altogether, these results suggest that pontine neurons connect to dFB neurons and regulate sleep via acetylcholine signals.

## Discussion

This study unravels post-synaptic neurons of the PB-FB pathway. We first focused on pontine neurons as assessed by anterograde tracing (Fig. 1a), and found that pontine neurons have dendrites in both the vFB and the dFB (Fig. 1c). We next found that P-FN neurons and pontine neurons have excitatory connections (Fig. 1d-h). We then investigated the role of pontine neurons in sleep regulation and identified that cholinergic pontine neurons promote arousal (Fig. 2). These results indicate that P-FN neurons activate pontine neurons, thus promoting arousal. However, we have not tested the functional impact of the connection between P-FN neurons and pontine neurons on sleep. Further study will clearly show the functional impact of the connection between the P-FN neurons and the pontine neurons on sleep regulation. Finally, we examined the relationship between pontine neurons and dFB neurons. As shown in Fig. 3a-c, we found that pontine neurons and dFB neurons form anatomical connections. We further investigated the specific neurons within FB interneurons that convey neuronal signals from P-FN neurons to dFB neurons to regulate arousal. Then we found that two types of FB interneurons named vDeltaB, C, D, and hDeltaC promote arousal (Fig. 3d). Although the functional connections between pontine neurons and dFB neurons are not demonstrated in this study, our results suggest that pontine neurons project to dFB neurons. In addition, knockdown of inhibitory acetylcholine receptor mAChR-B promotes sleep (Fig. 3e). This result suggests the possibility that acetylcholine signals from pontine neurons inhibit dFB neurons and promote arousal. Furthermore, a previous study showed that neurons that project to the ventral part of the FB (vFB neurons) promote sleep and mediate consolidation of long-term memory^31^. Since dendrites of pontine neurons arborize in the ventral part of the FB, there would be interactions between vFB neurons and pontine neurons. Further research will clarify the relationship between vFB neurons and pontine neurons in sleep and memory regulation.

According to previous reports, pontine neurons regulate optomotor behavior and express tachykinin, a neuropeptide that regulates aggression^20,22,32^. Additionally, T1 dopamine neurons, which are upstream of pontine neurons, regulate aggression as well^33^. Also, we recently showed that T1 dopamine neurons are upstream of P-FN neurons^11^. Besides, P2 neurons, which include FB interneurons, regulate chronic isolation evoked sleep loss^34^. Moreover, courtship-regulator P1 neurons activate T1 neurons and modulate sleep/courtship balance based on the nutritional status^35^. Taking the abovementioned into account, we consider that arousal signals related to aggression, courtship, nutrition, and vision converge into the PB-FB pathway and pontine neurons to regulate arousal. Further studies should clarify the mechanisms of these arousal signals in sleep regulation within the PB-FB pathway and pontine neurons.

In conclusion, our results unravel the functional connection of the PB-FB pathway to pontine neurons and the role of this circuit in sleep regulation. This circuit likely regulate dFB neurons via inhibitory acetylcholine signals. Taken together, our results offer a neuronal circuit basis for studying the mechanisms of sleep regulation (Fig. 3f).

## Acknowledgments

We thank Drs. Julie H. Simpson, Takaomi Sakai, Kristin Scott, Richard A. Baines, BDSC, VDRC, and DGRC for fly stocks, and the members of Kume lab for discussions. We also thank Dr. Takako Morimoto for providing insights into the pontine neurons.

## Author Contributions

YSK, JT, and KK designed the experiments. YSK conducted all experiments and data analysis. YSK wrote this manuscript and KK revised it.

