## Supplementary Figure S1 for "Interneurons of fan-shaped body promote arousal in *Drosophila*"

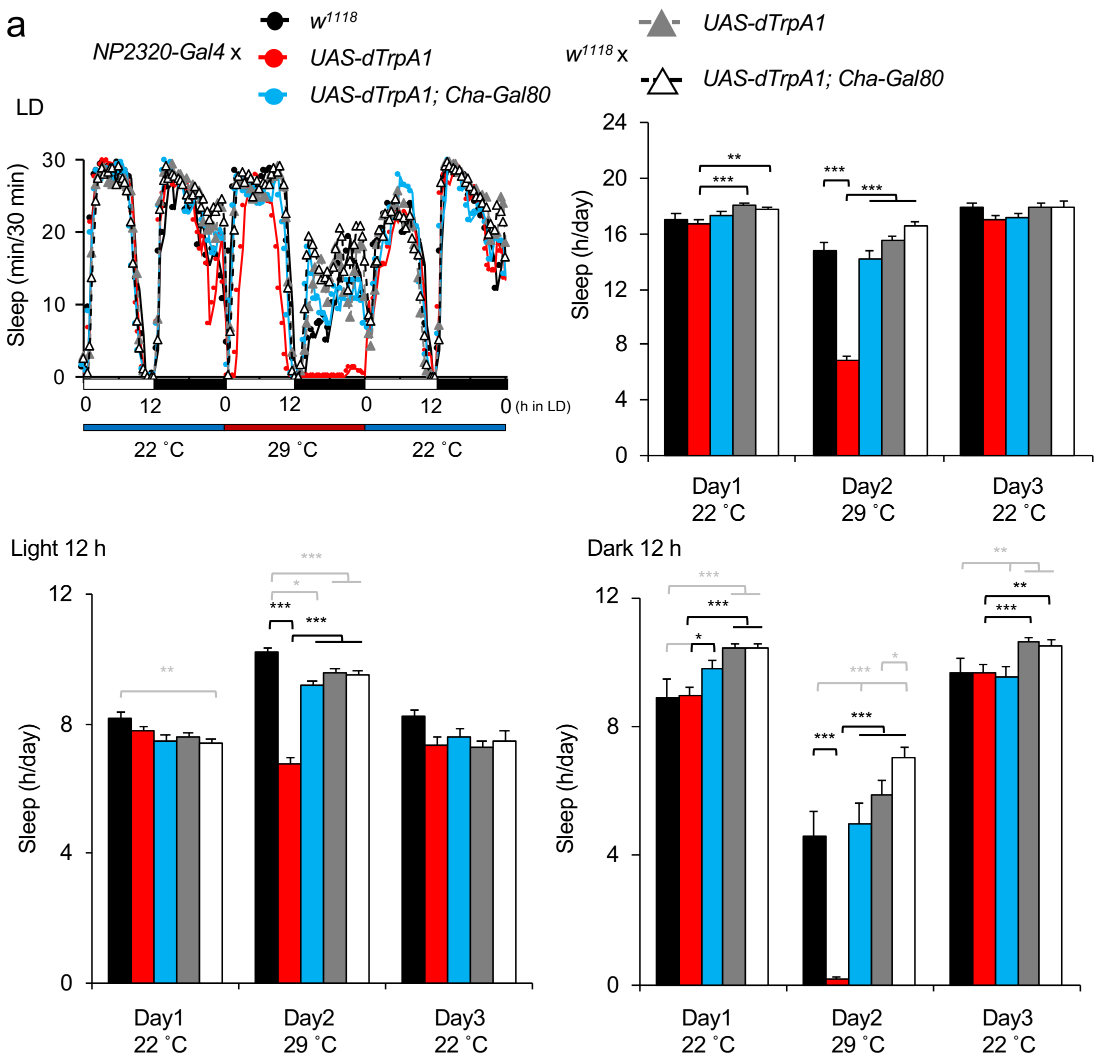


**Supplementary Figure S1.** Activation of the pontine neurons with *NP2320-Gal4* in LD conditions decreased sleep especially in the dark phase. (a) top right: Sleep profile of each genotype (n = 20, 32, 21, 32, 32 respectively). Thermo-genetic activation by *dTrpA1* occurs at 29 ˚C but not at 22 ˚C. top left: Quantification of the sleep time in 24h. bottom: Quantification of the sleep time of 12 h in light (left) or dark (right) phase. Data are presented as mean + SEM. one-way ANOVA with a Tukey-Kramer HSD test was used. * P < 0.05, ** P < 0.01, *** P < 0.001
