## Supplementary figures and images for "Interneurons of fan-shaped body promote arousal in *Drosophila*"

### Supplementary Movie S1

#### Slide 1
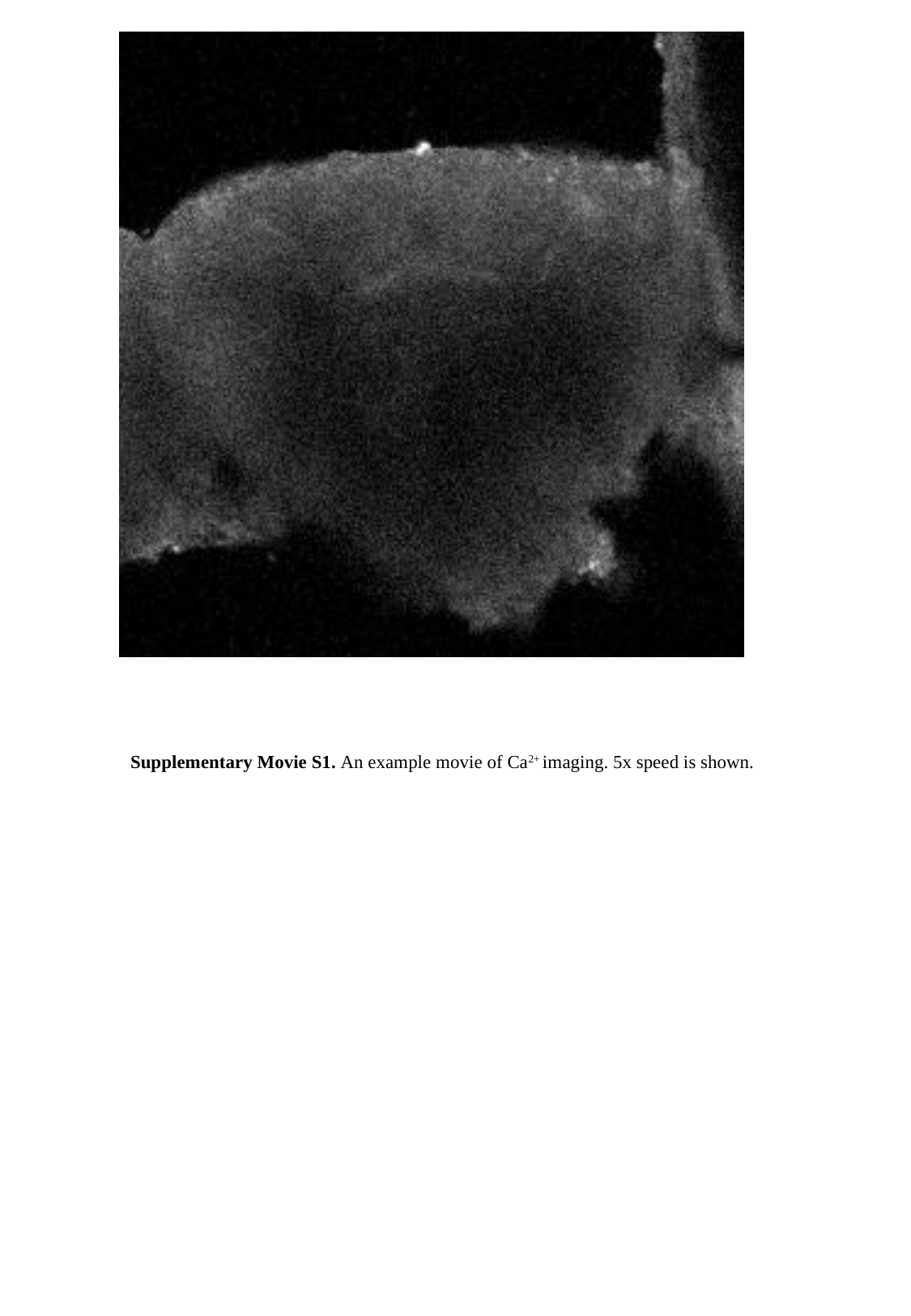

Supplementary Movie S1. An example movie of Ca^2+^ imaging. 5x speed is shown.
